# A radiofrequency coil for infants and toddlers

**DOI:** 10.1101/2022.10.17.512566

**Authors:** Kyle M. Gilbert, Emily S. Nichols, Joseph S. Gati, Emma G. Duerden

**Affiliations:** Centre for Functional and Metabolic Mapping, The University of Western Ontario, London, Ontario, Canada; Department of Medical Biophysics, The University of Western Ontario, London, Ontario, Canada; Applied Psychology, Faculty of Education, The University of Western Ontario, London, Ontario, Canada; Western Institute for Neuroscience, The University of Western Ontario, London, Ontario, Canada; Department of Pediatrics, Schulich School of Medicine and Dentistry, London, Ontario, Canada

**Keywords:** radiofrequency coil, paediatric imaging, motion correction, parallel imaging, neuroimaging, magnetic resonance imaging

## Abstract

**Background:** Infants and toddlers are a challenging population on which to perform MRI of the brain, both in research and clinical settings. Due to the large range in head size during the early years of development, paediatric neuro-MRI requires a radiofrequency (RF) coil, or set of coils, that is tailored to head size to provide the highest image quality. Mitigating techniques must also be employed to reduce and correct for subject motion.

**Objective:** To develop an RF coil with a tailored mechanical-electrical design that can adapt to the head size of three-month-old infants to three-year-old toddlers.

**Materials and methods:** An RF coil was designed with tight-fitting coil elements to improve SNR in comparison to commercially available adult head coils, while simultaneously aiding in immobilization. The coil was designed without visual obstruction to facilitate an unimpeded view of the child’s face and the potential application of camera or motion-tracking systems.

**Results:** Despite the lack of elements over the face, the paediatric coil produced higher SNR over most of the brain compared to adult coils, including more than 2-fold in the periphery. Acceleration rates of 4-fold in each Cartesian direction could be achieved. Higher SNR allowed for shorter acquisition times through accelerated imaging protocols, reducing the probability of motion during a scan.

**Conclusion:** Modification to the acquisition protocol, with immobilization of the head through the adjustable coil geometry, and subsequently combined with a motion tracking system provides a compelling platform for scanning paediatric populations without sedation and with improved image quality.

## 1 Introduction

Infant and toddler populations are a challenging cohort for clinical imaging due to their inability to remain still during an MRI exam. This difficulty is confounded with the relatively few vendor-supplied and commercial options for radiofrequency (RF) coils tailored to infant and toddler populations as compared to those available for adult imaging. When designed with the specific challenges of young paediatric neuroimaging in mind, the RF coil can aid in mitigating motion and improving image quality.

Two of the primary technical challenges when developing an RF coil for paediatric MRI of the brain are (i) retaining high coil sensitivity (and therefore image SNR) over a range of head sizes and (ii) mitigating head motion. Coil elements must be placed proximal to the subject to produce the highest sensitivity; however, this can be difficult to accomplish over the entire age range due to the variance in head size. Coils tailored to the paediatric age group of interest have been developed to address this problem[1], including an entire suite of paediatric coils optimized for multiple age groups[2,3]. To negate the need for multiple coils, size-adaptable designs have been developed that can provide a tight fit over a range of head sizes[4–6].

Head motion is the second ubiquitous challenge when imaging these age groups. Motion can lead to image blurring, ghosting, and distortions[7], and is exacerbated in younger paediatric populations due to their predilection for moving when subjected to the stressors of being in an MRI scanner while awake. MRI scans are often acquired during natural sleep; however, infants may still move during sleep or awaken during sequences. In the clinical setting, sedation and anesthesia are often administered to address motion, albeit with intrinsic risk to the patient.

Even under the influence of anesthesia, patient motion may persist, with the dominant direction of movement being toward the foot end of the bed[8]; awake children also demonstrate a significant nodding movement[8]. Tailoring the coil housing to the shape of the participant’s head, as well as employing an aforementioned adaptable coil design, can aid in mitigating this motion. These strategies become even more effective when combined with pulse-sequence optimizations that reduce scan times[9], such as parallel imaging[10,11] and compressed sensing[12,13]—these techniques benefit substantially from the increased SNR afforded by dedicated paediatric coils. For younger infants, feeding and swaddling can promote sleep to effectively reduce motion[14]. Nonetheless, residual motion persists—notably for older infants and particularly awake toddlers— that can degrade image quality and produce unusable images. Prospective and retrospective motion correction has therefore been introduced to address this residual motion. By tracking motion in real-time using navigators[15] or external hardware[16–18], translational and rotational estimates can be used to modify RF and gradient pulses to ensure alignment between the head position and imaging volume (i.e., prospective motion correction) or correct k-space trajectories during image reconstruction (i.e., retrospective motion correction)[19].

This manuscript presents a tailored solution for imaging infants and toddlers aged three months to three years on a 3T MRI scanner. The children were scanned, either sleeping or awake, without the use of and potential risks of anesthesia. A mechanically conforming RF coil was designed that can adapt to the range in head size of the infant and toddler age groups to simultaneously increase SNR and help immobilize the head: this adaptive topology was modelled after the design presented by Ghotra et al.[5] (for infants aged 1 – 18 months) with important modifications made to accommodate a wider age range. All computer-aided design (CAD) files have been provided to add this solution to the limited repertoire of publicly available paediatric coil designs. An open-face configuration was employed to accommodate the potential application of a camera or motion-tracking system—the consequences of which are evaluated in comparison to conventional adult coils.

## 2 Materials and methods

### 2.1 Radiofrequency coil

#### 2.1.1 Coil mechanicals

The coil housing was designed to accommodate children between the ages of three months and three years. The coil was split into four disparate sections to adapt to the variability in head size within the age range (Figure 1). The geometry for the inner surface of the housing was based off the average head shape of a 3-year old (*n* = 36; provided by the Neurodevelopmental MRI Database[20,21]) and allowed for approximately 6 mm of space for foam cushioning. The arch of the neck was included in the coil housing to allow closer proximity of coils and to create a “cup” for the occiput of the head to provide better immobilization.

**Fig. 1.**
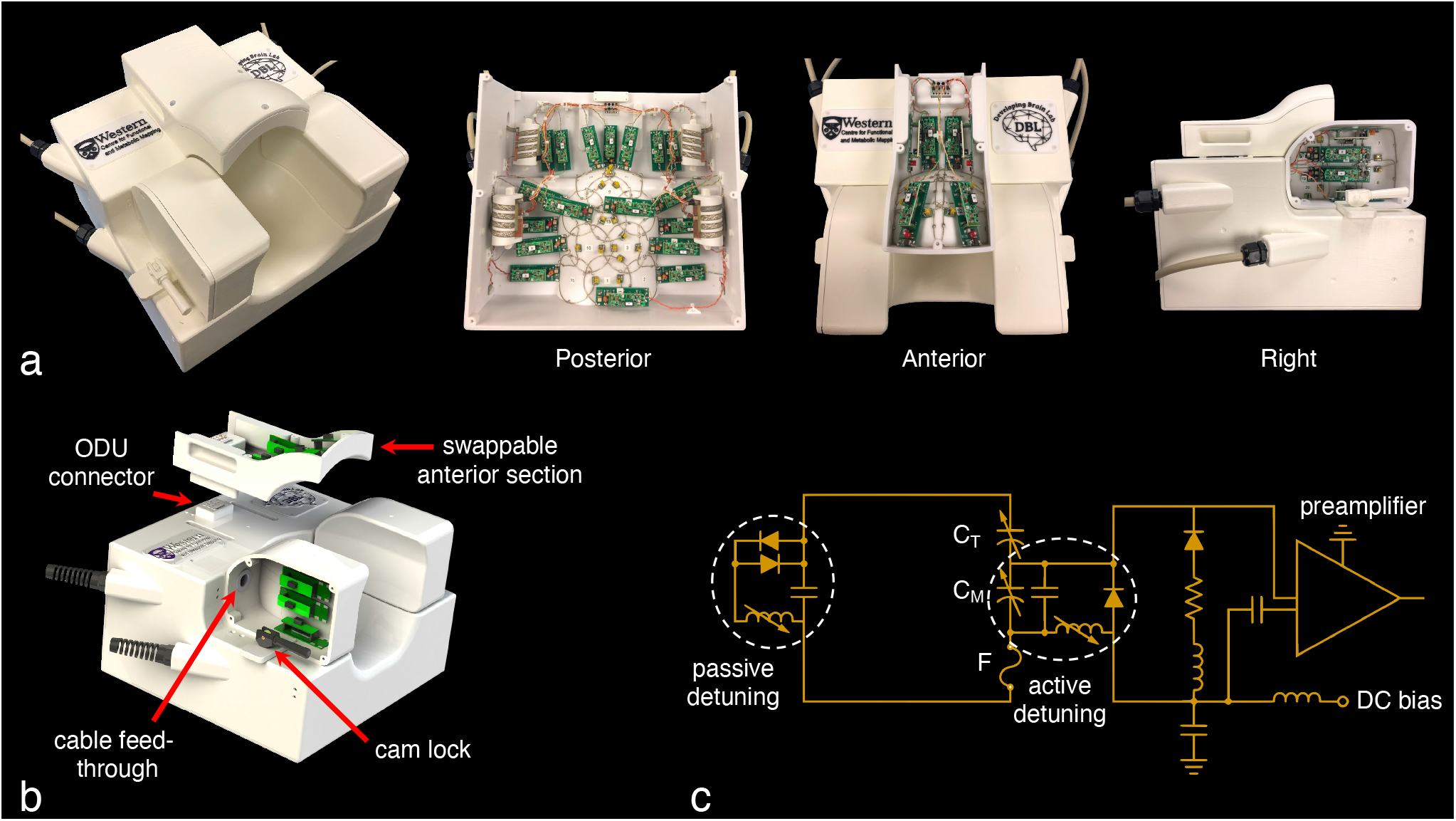
**a** Photographs of the 32-channel paediatric coil. **b** Lateral sections can slide laterally to accommodate varying head width and are fixated afterwards with custom cam locks. The coaxial cables of lateral elements are routed through the housing to the posterior section. Three interchangeable anterior sections were constructed to accommodate the large range in head size over the desired age range (three months to three years old). High-density ODU connectors connect coaxial and DC cables from the anterior to posterior sections, precluding the need for a cumbersome plug on the anterior sections. **c** The circuit schematic of a single receive element depicts the active and passive detuning networks. Coil elements are connected directly to low-input-impedance preamplifiers. C_T_: tuning capacitor; C_M_: matching capacitor; F: fuse.

The large posterior section of the design was stationary, while the left and right sections could slide laterally to create a housing width anywhere between 12.0 cm and 17.1 cm (excluding space for foam). Once the lateral sections were positioned as close as possible to the subject’s head, they could be locked into place using custom-built cam locks. This served two purposes: (i) to increase coil sensitivity by ensuring the closest possible proximity to the brain and (ii) to mitigate head motion. This adaptive topology was based off of the design presented by Ghotra et al.[5] (for infants aged 1 – 18 months); however, the older and wider age range desired for this study necessitated design modifications to accommodate a larger variance in the anterior-posterior dimension of the head.

The variance in head size in the anterior-posterior direction over the desired age range was addressed by constructing three interchangeable anterior sections based off the average head sizes of a three-month old, one-year old, and three-year old[20,21]. The anterior-posterior dimension of the housing was therefore either 16.5 cm, 18.5 cm, or 20.0 cm (Figure 2). No coil elements or mechanical structure were placed in front of the subject’s face, inferior to the brow ridge, to accommodate the use of a camera or motion-tracking system.

**Fig. 2.**
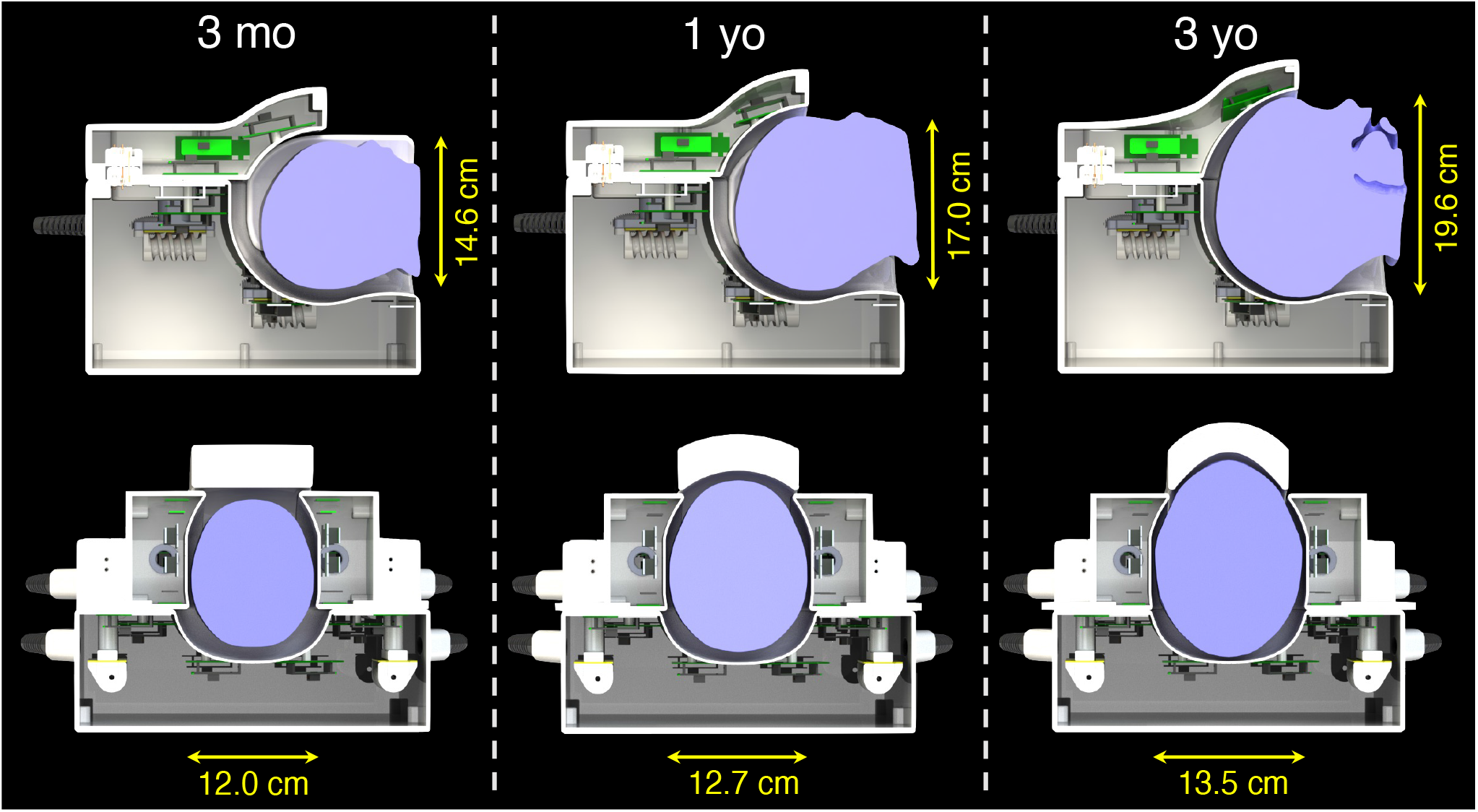
Sagittal and axial cross-sections of the three coil configurations: for children less than three months of age, between three months and one year of age, and between one and three years of age. The representative heads approximate the average size of a three-month old, one-year old, and three-year old[20,21], with dimensions provided along the anterior-posterior and left-right directions. The adjustability of the lateral and anterior sections allows for consistent loading across the age range.

All CAD drawings have been made openly available to facilitate reproducibility: https://doi.org/10.17605/OSF.IO/6EZ4Q.

#### 2.1.2 Coil circuitry

Each receive-coil configuration (i.e., using one of three possible anterior sections) was comprised of 32 loop elements arranged in a soccer-ball geometry[22] with adjacent elements within each section overlapped to reduce inductive coupling[23]. Fourteen elements were placed on the posterior section, 6 elements on the anterior section, and 6 elements on each of the right and left sections. Elements were tuned to 123.2 MHz and matched to 200 + j50 Ω (i.e., the optimal impedance to minimize the noise figure of preamplifiers) when loaded with a head-shaped phantom of a 2-year-old filled with 50-mM sodium chloride and water. A parallel-resonant, active-detuning circuit with a high-power PIN diode (Macom MA4P7435NM-1091T) was placed near the input of each receive element. A parallel-resonant, passive-detuning circuit[24] was added to each element as a secondary detuning method and to enhance detuning during transmission. As an extra precaution, a fast-blow fuse (315-mA limit) was added to the daughterboard of the preamplifier. A circuit schematic of a receive element is provided in Figure 1c.

Most preamplifiers were located directly at the input of their respective coil element. Preamplifiers (Stark Contrast, Erlangen, Germany) had a low input impedance to facilitate preamplifier decoupling[23] and an integrated cable trap between the first and second amplification stage. Two elements on each lateral section were connected to their respective preamplifier, located in the posterior section of the housing, through 8.9-cm-long flexible coaxial cables. Coaxial cables from the lateral elements were routed to the posterior section through a feed-through that would move with the lateral housing. Coaxial cables from the anterior sections were routed to the posterior section through high-density coaxial connectors built into the coil housing (ODU-MAC product line). From the posterior section, each multi-conductor cable was wound into a large choke balun before exiting the housing—this prevented common-mode currents on the coaxial-cable grounding shields. Preamplifier decoupling and active detuning of each receive element were measured using standard double-probe techniques described by Keil et al.[2]. The coupling between overlapping receive elements, *S*_ij_, was measured from the preamplifier inputs.

### 2.2 Imaging

All imaging was performed at the Centre for Functional and Metabolic Mapping at the University of Western Ontario. MRI data collection was performed on a human, whole-body Siemens Prisma Fit scanner (Siemens Healthineers AG, Erlangen, Germany) operating with a 3-T main magnetic field and an XR80/200 gradient coil (maximum gradient strength: 80 mT/m, maximum slew rate: 200 T/m/s). The system is equipped with 128 receiver channels, of which 32 were utilized: plugs were interfaced to Tim coil interface legacy adaptors (part numbers: 14460282 and 14418519) to allow operation on the Siemens Prisma hardware platform.

All human imaging was performed in accordance with the human subject research ethics board (HSREB) at the University of Western Ontario (REB #115440). All imaging was performed with the informed consent of a parent or legal guardian. Rigorous testing of both coil and patient safety was performed before the scanning of human subjects to ensure robust operation and subject safety. Testing followed regulatory guidelines specified by the IEC[25,26], summarized by Hoffmann et al.[27], and described in a previous study by the authors[28].

### 2.3 Coil performance

The coil performance was evaluated by assessing image SNR, noise level, noise correlation, and geometry factor. The performance was compared to two vendor-supplied commercial coils intended for adult imaging (Siemens Healthineers AG, Erlangen, Germany): (i) a 20-channel head and neck coil (routinely employed for paediatric neuroimaging within our facility due to its large dimensions (approximately 26.8 cm (anterior-posterior) by 22.8 cm (left-right)) that can accommodate an infant’s shoulders, and (ii) a 32-channel head-only coil (approximate dimensions: 23.0 cm (anterior-posterior) by 19.6 cm (left-right))—chosen due to it having the same number of coil elements as the paediatric coil.

Performance metrics were acquired with a phantom that approximated the size of a 2-year-old’s head and was filled with 50-mM sodium chloride and water. The largest anterior section was required to accommodate the head phantom. Image SNR maps were derived from gradient-echo images, with and without RF transmission, using a sum-of-squares reconstruction[23,29] (matrix size: 96 × 96, field of view: 192 × 192 mm, number of slices: 96, slice thickness: 2 mm, TE/TR: 5/875 ms, flip angle: 20°, bandwidth: 250 Hz/pixel, number of averages: 4). The noise-only acquisition was additionally used to calculate the noise correlation matrix and the normalized noise level of each channel. Geometry-factor maps were created by retrospectively under-sampling the fully sampled k-space data and reconstructing using SENSE[11]. All calculations were performed in Matlab (MathWorks, Natick, MA, USA).

To assess the benefits of having adjustable lateral sections and multiple anterior sections, the SNR was measured of a phantom that approximated the head size of a 3-month-old. The SNR was compared when the coil was placed in its largest configuration (i.e., lateral sections as wide as possible and using the largest anterior section) versus when in its smallest configuration (i.e., lateral elements placed tight to the phantom and using the smallest anterior section).

Image quality was assessed by scanning a three-month-old infant in the coil’s smallest configuration. A turbo-spin-echo sequence was employed with a total acquisition time of 2 min 33 s (matrix size: 192 × 156, field of view: 192 × 156 mm, number of slices: 115, slice thickness: 1 mm, TE/TR: 140/6150 ms, refocusing flip angle: 120°, echo-train length: 19, bandwidth: 190 Hz/pixel).

## 3 Results

### 3.1 Coil performance

Active detuning provided a minimum isolation of 34 dB during transmission. The reference voltage required by the scanner’s body transmit coil, measured with and without the receive coil present, differed by less than 2.5% (for all three coil configurations—i.e., different anterior sections). Actual flip-angle maps showed no regions of increased flip-angle due to the presence of the receive coil.

RF and gradient heating tests produced negligible changes in the temperature of the coil housing and components, with less than a 3% reduction in image SNR post-heating.

The mean geometric decoupling between adjacent elements was -16 dB (worst-case: -10 dB). Preamplifier decoupling had a mean of -26 dB across all elements (worst-case: -20 dB). The mean and maximum noise correlation was 9.5% and 59%, respectively (Figure 3). In contrast, the adult head/neck and head-only coils had higher mean noise correlations (17% and 25%, respectively) due to being insufficiently loaded by the “two-year-old” phantom. The normalized noise level of individual channels ranged from 0.76 – 1.46-fold, compared to 0.81 – 1.29-fold and 0.71 – 1.67-fold for the adult head/neck and head-only coils, respectively.

**Fig. 3.**
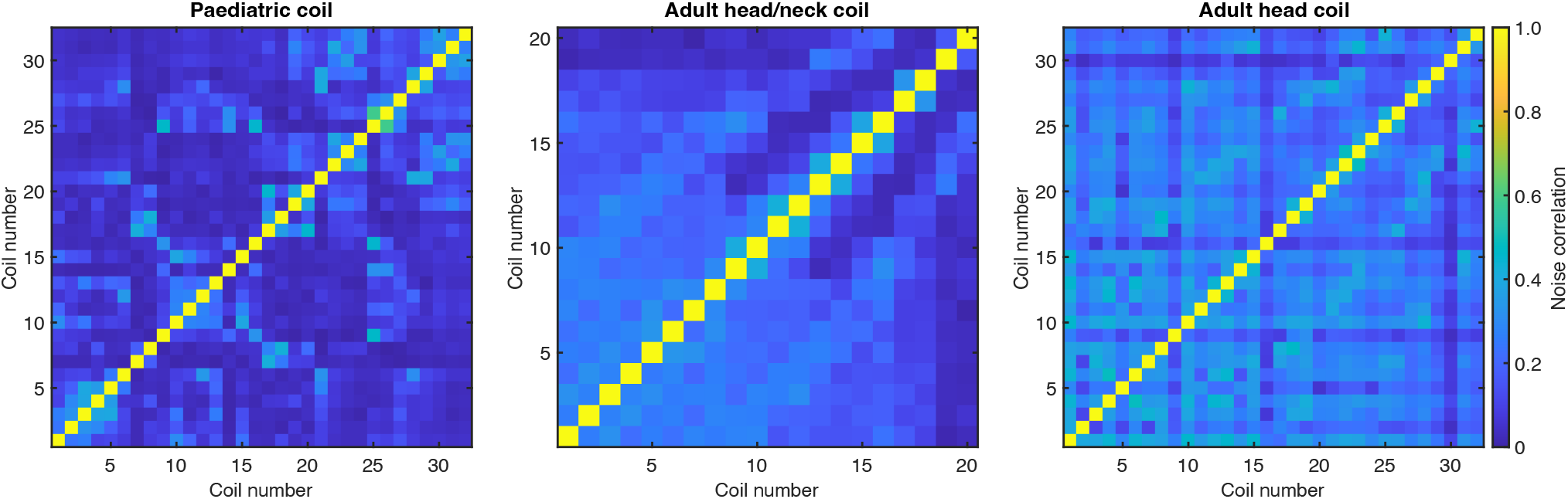
Noise-correlation matrices of the paediatric coil, 20-channel adult head/neck coil, and 32-channel adult head-only coil. The paediatric coil produces a lower mean noise correlation (9.5% compared to 17%/25% for the adult head/neck and head-only coils, respectively), due to being optimized for a smaller load.

The paediatric coil produced higher SNR throughout the majority of the brain relative to the adult coils, in large part due to the smaller dimensions of the paediatric coil (Figure 4). The relative improvement between the coils was spatially varying (see Table 1), with the largest gains (>2-fold) in the periphery (cortex) of the brain. Superior, frontal, and lateral regions demonstrated relative improvements between 1.14 – 1.98-fold. In the paediatric coil, there were no coil elements placed in front of the face (unlike for the larger adult head/neck coil)—this design topology was adopted to allow a motion-tracking camera to view the entire face, an otherwise difficult task to accomplish when using a small, tight-fitting coil; however, this also resulted in lower SNR in the centre of the brain (0.73-fold). In contrast, when compared to the adult head coil—which has fewer receive elements above the face—the paediatric coil produced a 1.04-fold SNR improvement in the central region.

**Table 1.** SNR ratio between the paediatric coil and the adult head/neck and head-only coils for ROIs illustrated in Figure 4

| ROI | SNR<sub>paediatric</sub> / SNR<sub>adult</sub> |  |
| --- | --- | --- |
|  | Adult head/neck coil | Adult paediatric coil |
| central | 0.73 | 1.04 |
| frontal | 1.14 | 1.40 |
| peripheral | 2.00 | 2.32 |
| lateral | 1.34 | 1.58 |
| superior | 1.89 | 1.98 |

**Fig. 4.**
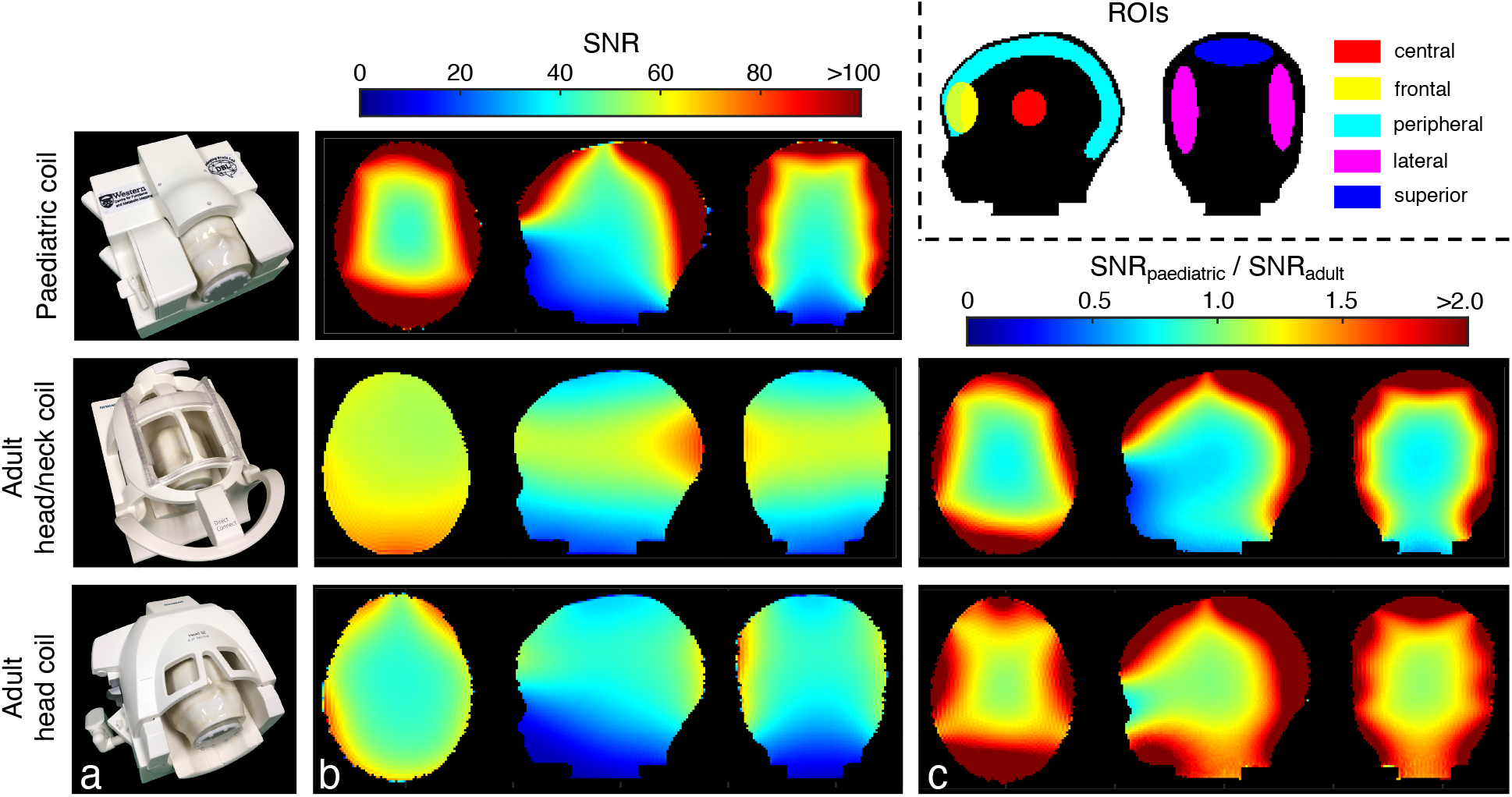
**a** Photographs of the “two-year-old” head phantom within the respective coils depicts the realistic positioning of a two-year-old subject. **b** The image SNR of the head phantom, as acquired with the paediatric coil, adult head/neck coil, and adult head-only coils, shows spatially varying SNR, mainly attributable to the difference in coil size. **c** The resultant ratio between image SNR attained with the paediatric coil versus the adult coils demonstrates large increases in the periphery. The inset provides the ROIs over which the mean SNR was calculated (see Table 1).

The SNR of a “3-month-old” head phantom when the coil was in its tightest configuration versus when in its largest configuration showed a 1.71-fold increase in the frontal region, 1.69-fold increase in the lateral regions, and a 1.15-fold increase in the centre of the head (Figure 5). Negligible differences were present in the posterior region of the coil, as the posterior section is non-adjustable.

**Fig. 5.**
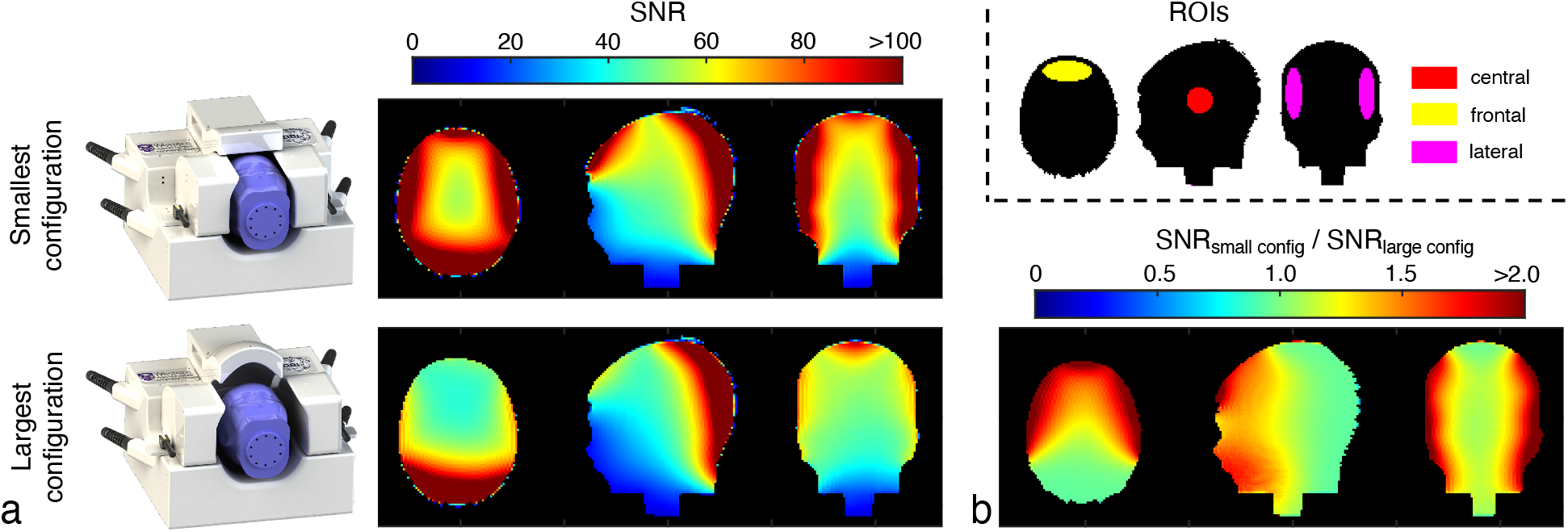
**a** SNR of a “three-month-old” head phantom when in the smallest coil configuration and when in the largest coil configuration. **b** The SNR ratio between the two configurations shows a 1.71-fold increase in the frontal region, 1.69-fold increase in the lateral regions, and 1.15-fold increase in the central region when employing the tightest coil configuration. The inset provides the ROIs over which SNR ratios were calculated.

The paediatric coil produced substantially lower geometry factors across all acceleration rates compared to the adult coils (Figure 6). The paediatric coil could achieve acceleration rates of three-or four-fold along each Cartesian direction before a substantial degradation in accelerated SNR would occur.

**Fig. 6.**
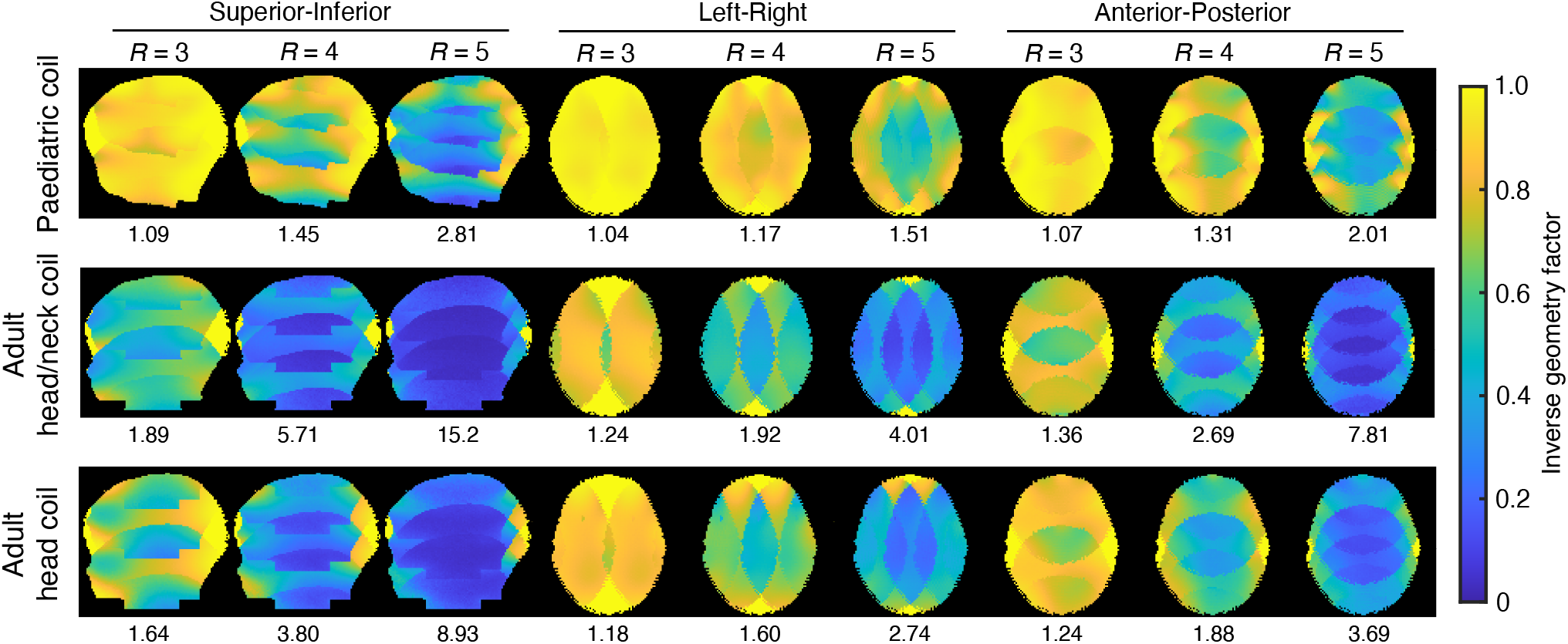
Inverse geometry factor maps for the 32-channel paediatric coil, 20-channel adult head/neck coil, and 32-channel adult head-only coil for acceleration rates of three, four, and five along the superior-inferior, left-right, and anterior-posterior directions. The mean geometry factor is provided beneath each respective map. The paediatric coil demonstrates markedly lower geometry factors across all acceleration rates due to its smaller geometry and therefore more spatially independent receive profiles.

The resultant quality of a turbo-spin-echo image acquired of a three-month-old infant is provided in Figure 7. Although motion between participants in this age range is highly dependent on their ability to sleep during the scan, no motion artifacts (ghosting or blurring) were present in this image.

**Fig. 7.**
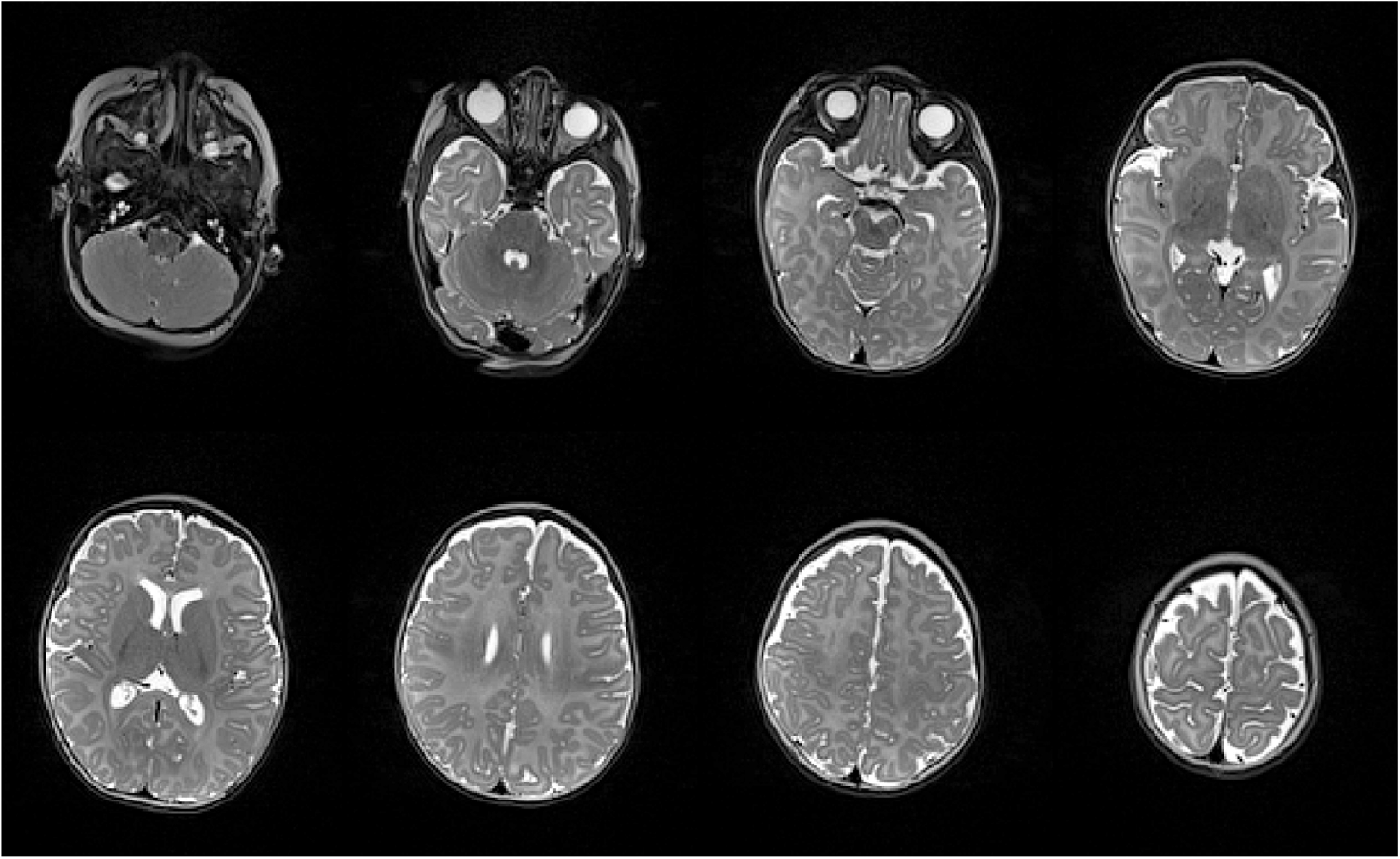
Representative axial slices of a turbo-spin-echo image acquired of a three-month-old infant when in the smallest coil configuration. No motion artifacts (ghosting or blurring) are visible in the image.

## 4 Discussion

The paediatric coil was designed with no elements or housing in front of the face (below the brow ridge) to accommodate the use of a camera or motion-tracking system that requires a clear view of the subject’s face while inside the bore of the MRI scanner[17]. Despite this design constraint, the paediatric coil produced superior SNR throughout most of the head compared to two vendor-provided adult coils—this can be primarily attributed to the smaller coil elements and the proximity of these coils to the head through an adjustable design. Despite the large SNR gains in the periphery of the brain (greater than 2-fold), a relatively small decrease in SNR was observed in the deeper regions of the head relative to the adult head/neck coil. This was in part due to the lack of coil elements in front of the face; however, this was deemed an acceptable compromise to facilitate the necessary monitoring of the child using cameras and/or motion tracking systems that could alleviate large decreases in SNR caused by subject motion. The SNR ratio to the adult head-only coil was generally consistent with gains reported by Ghotra et al.[5] when compared to a dedicated neonatal coil (for children aged 1 – 18 months).

The SNR comparison in this study was performed on a phantom equivalent to the size of a two-year-old’s head—this was to determine the lower-bound in potential SNR gains of the paediatric coil. Since the paediatric coil is adjustable in size, SNR gains are expected to be even higher when imaging smaller head sizes compared to the fixed geometry of conventional adult coils.

Scans of the “3-month-old” head phantom in the coil’s tightest configuration produced a substantial improvement in SNR in the frontal region (1.71-fold) and lateral regions (1.69-fold) compared to scanning in the coil’s largest configuration. This demonstrates the importance of being able to substantially vary the coil size in both the left-right and anterior-posterior dimensions—not just in comparison to an adult but over the 3-month (infant) to 3-year (toddler) age range.

The paediatric coil produced superior geometry factors compared to the adult coils, despite the open-face design. This is due to the closer proximity of its coil elements and therefore the higher spatial independence of the commensurate sensitivity profiles. When combined with higher SNR (as described above), the result is the ability to employ higher acceleration rates to reduce scanning time, thus reducing the probability of motion occurring during the acquisition.

## 5 Conclusion

Modification to the acquisition protocol, with immobilization of the head through the adjustable coil geometry, and subsequently combined with a motion tracking system produces a compelling platform for scanning paediatric populations without sedation and with improved image quality.

## Data availability

The data that support the findings of this study are openly available in an OSF repository (DOI: 10.17605/OSF.IO/6EZ4Q). This includes CAD files of the RF coil and performance data.

## Acknowledgments

The authors would like to thank Peter Zeman for fabrication assistance. This work was supported by the Canada Foundation for Innovation; Canada First Research Excellence Fund to BrainsCAN; Brain Canada Platform Support Grant; the Molly Towell Perinatal Research Foundation.

